## Supplementary material for "Can fMRI reveal the representation of syntactic structure in the brain?": Detailed example illustrating complete and incomplete subtrees

The figure below shows a constituency tree generated using the self-attentive encoder-based parser [25] for the sentence “Harry had never believed he would meet a boy he hated more than Dudley, but that was before he met Draco Malfoy.” This is the first sentence of the text presented to the subjects.

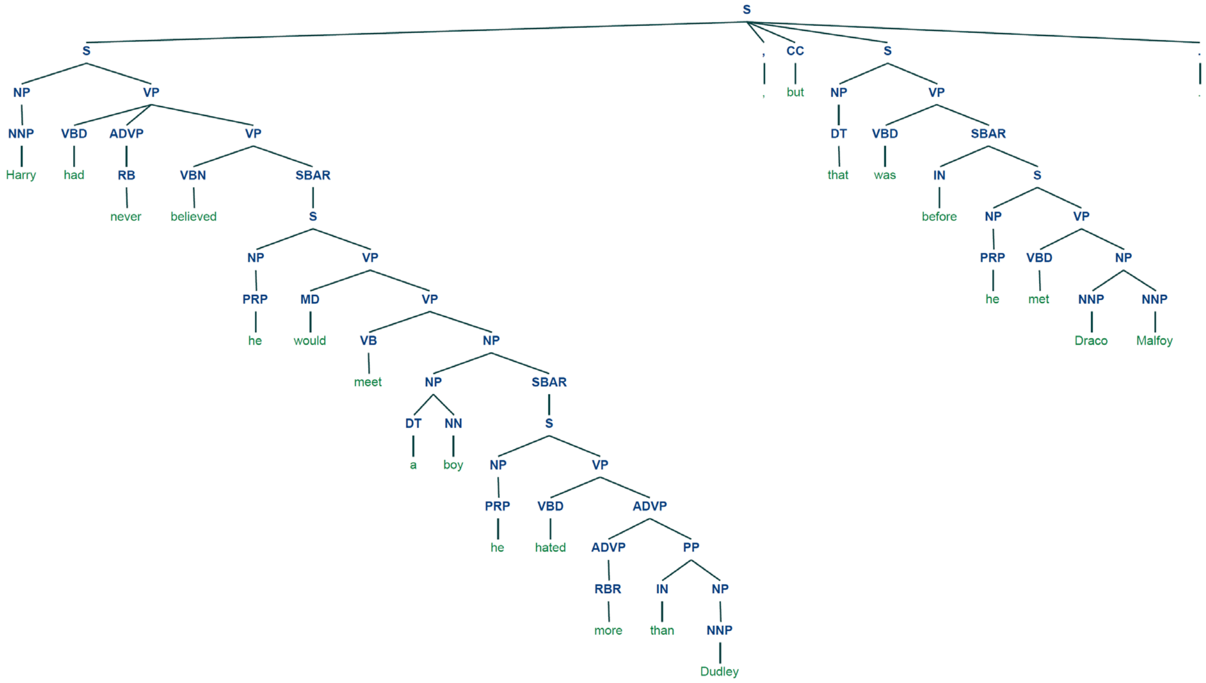


### Examples of complete subtrees used to generate the ConTreGE Comp vectors

We generate a ConTreGE Comp vector for every word using the largest subtree completed by that word. A subtree is considered complete when all of its leaves are terminals. The largest subtree completed by a given word refers to the subtree with the largest height that also satisfies the following conditions:

1. The given word must be one of its leaves
2. All of its leaves must only contain words that have been seen till then.

We now present the largest subtrees completed by a few of the words in the above sentence:

1. Harry

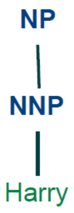

2. believed

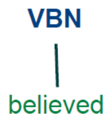


3. boy

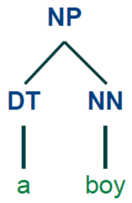


4. Dudley

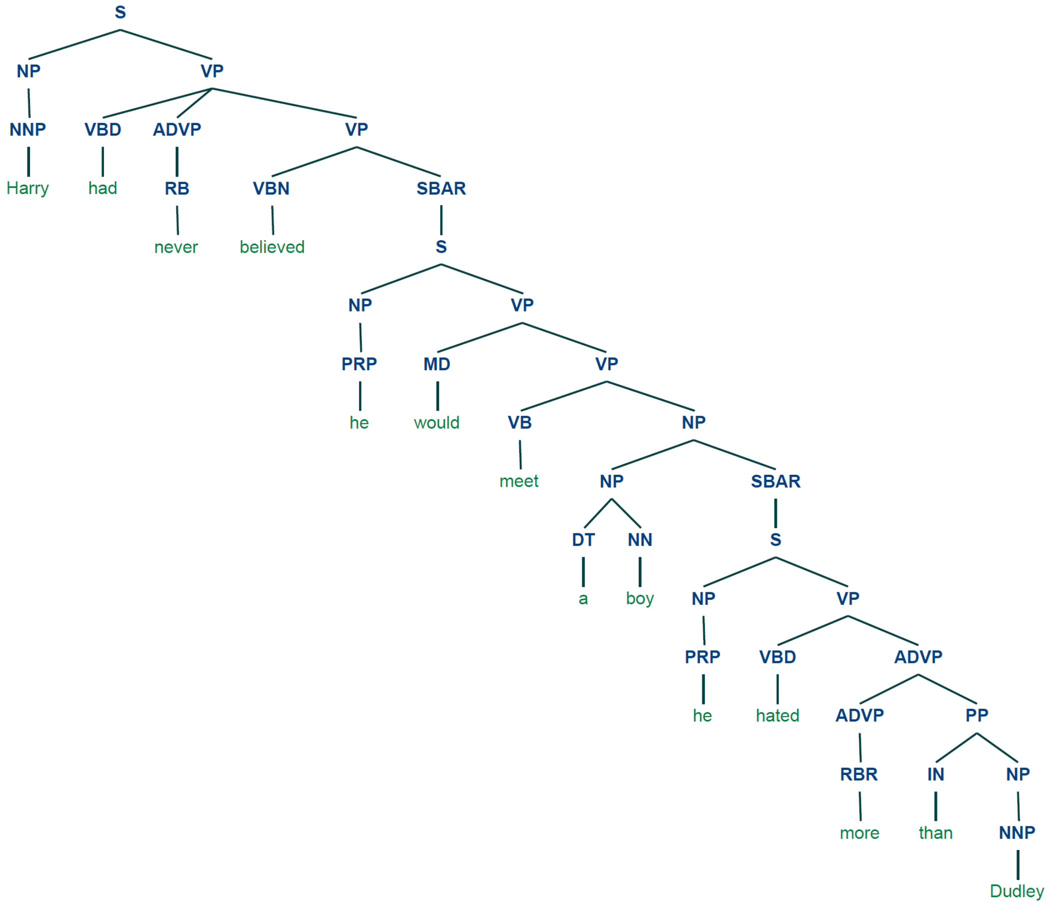


5. but

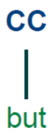

6. Draco

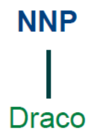


7. Malfoy

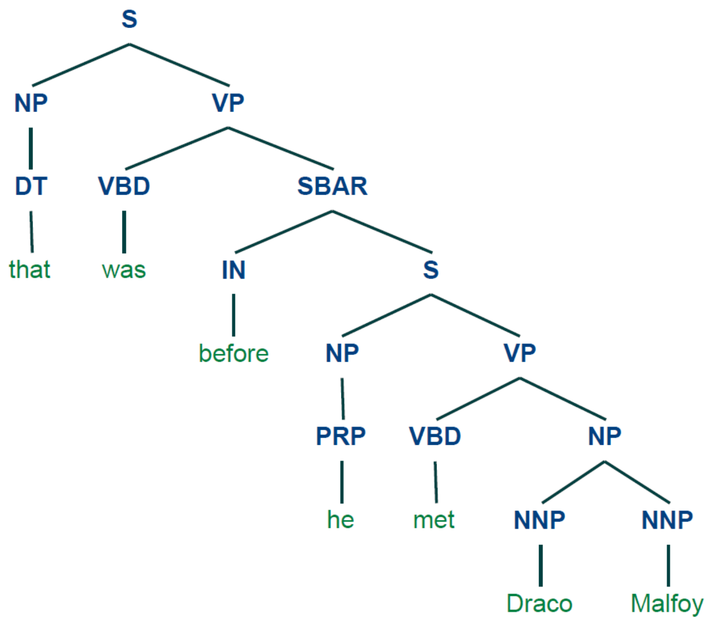


Each of these trees is encoded using the procedure described in the main paper to obtain one embedding per word.

### Examples of incomplete subtrees used to generate the ConTreGE Incomp vectors

We generate a ConTreGE Incomp vector for every word using the incomplete subtree that contains all of the Phrase Structure Grammar (PSG) productions needed to derive the words seen till then, starting from the root of the sentence's tree. A non-terminal node is expanded only if it eventually leads to the derivation of a word that has been seen. Any other non-terminal nodes are not expanded.

We now present the incomplete subtrees for a few of the words in the aforementioned sentence:

1. Harry (words seen till then = Harry)

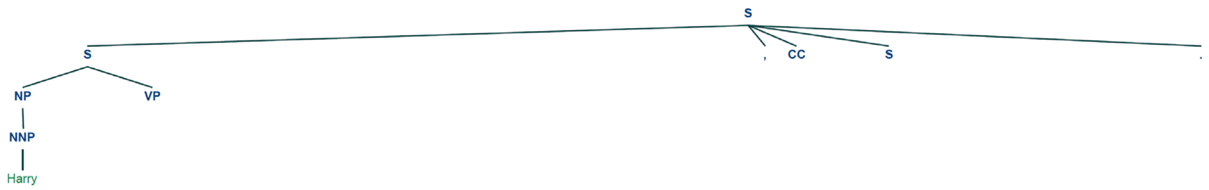

2. believed (words seen till then = Harry had never believed)

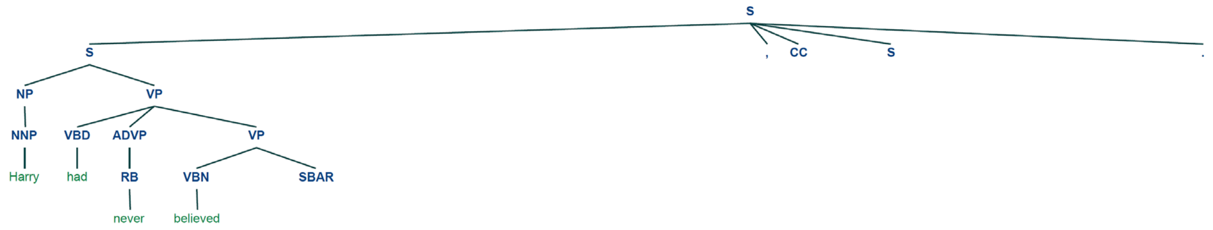


3. boy (words seen till then = Harry had never believed he would meet a boy)

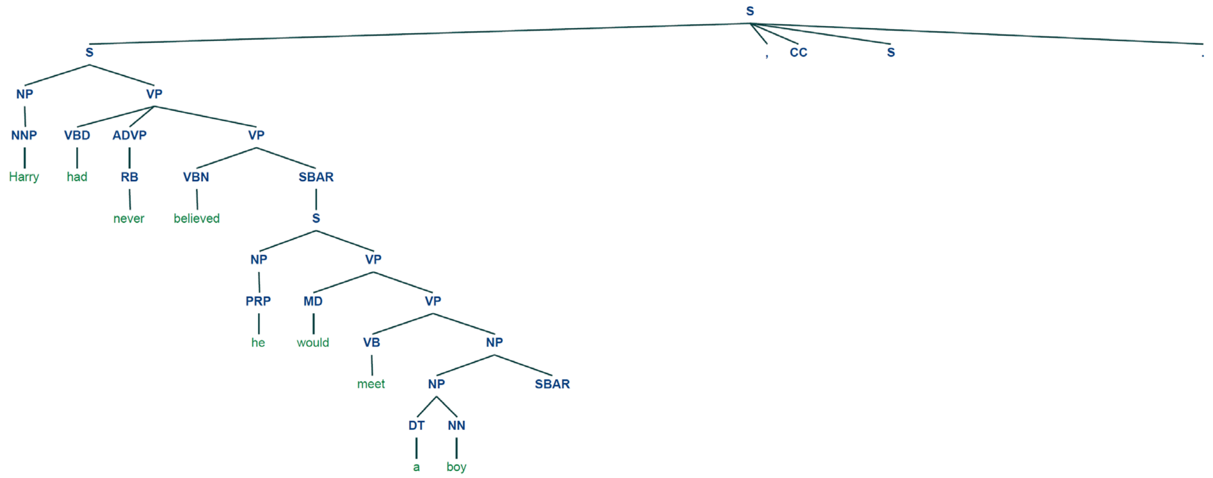


4. Dudley (words seen till then = Harry had never believed he would meet a boy he hated more than Dudley)

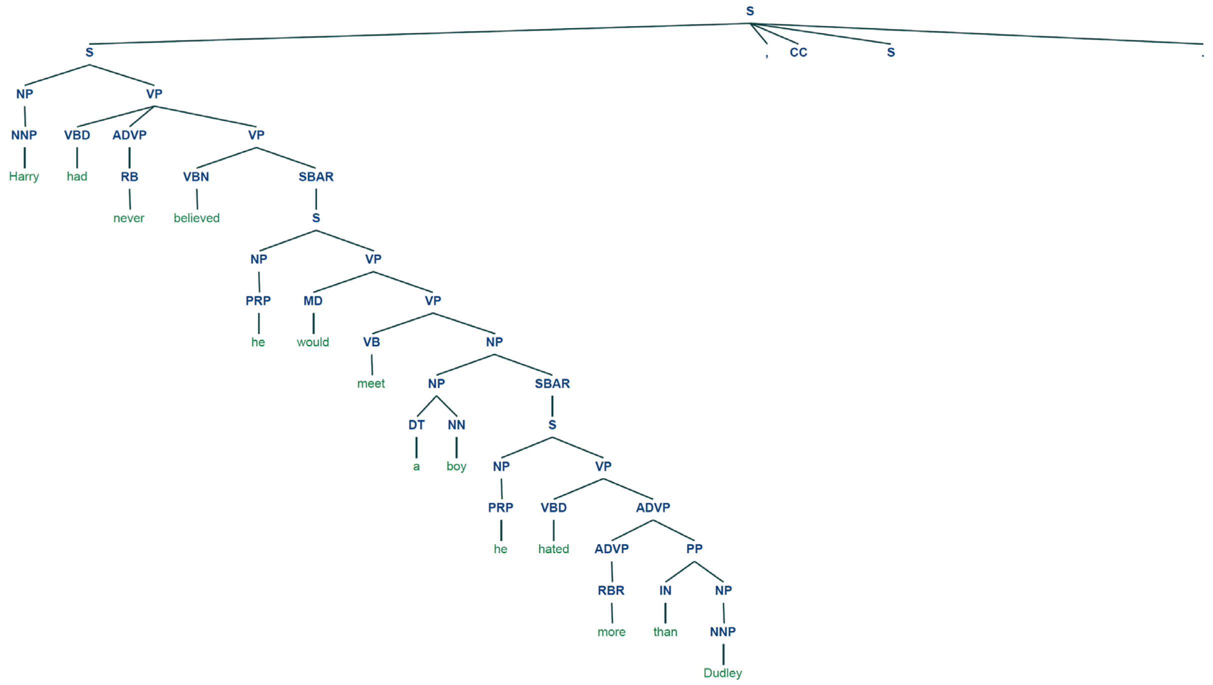

5. but (words seen till then = Harry had never believed he would meet a boy he hated more than Dudley, but)

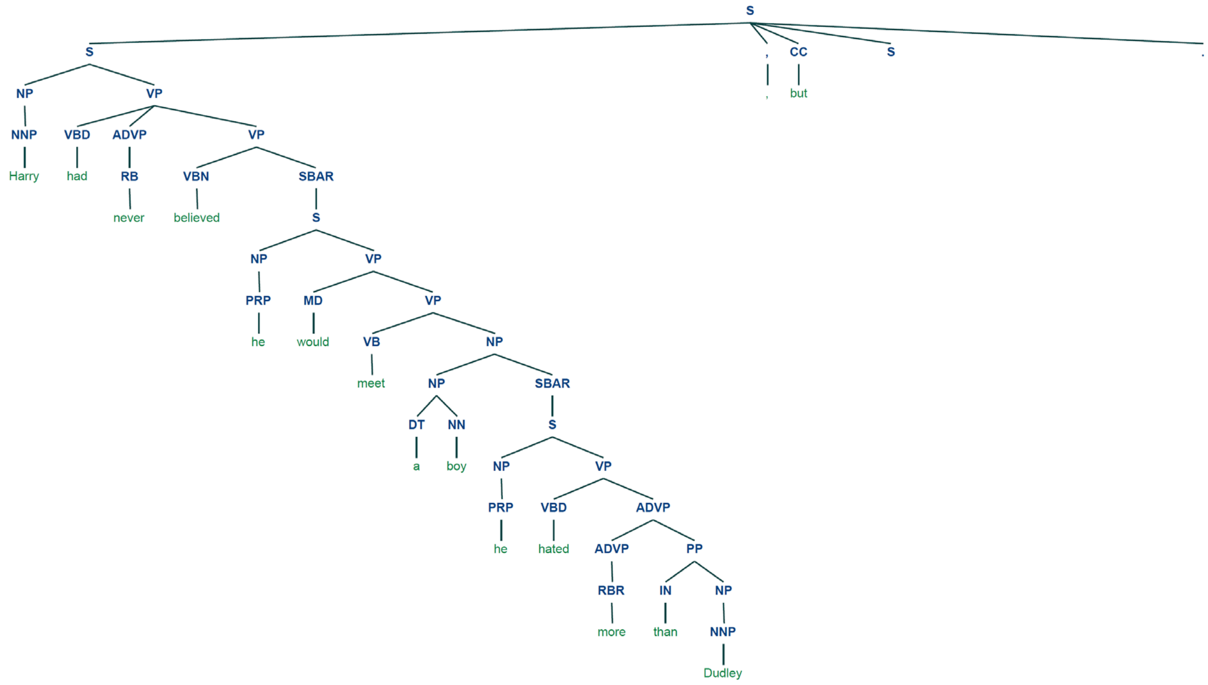


6. Draco (words seen till then = Harry had never believed he would meet a boy he hated more than Dudley, but that was before he met Draco)

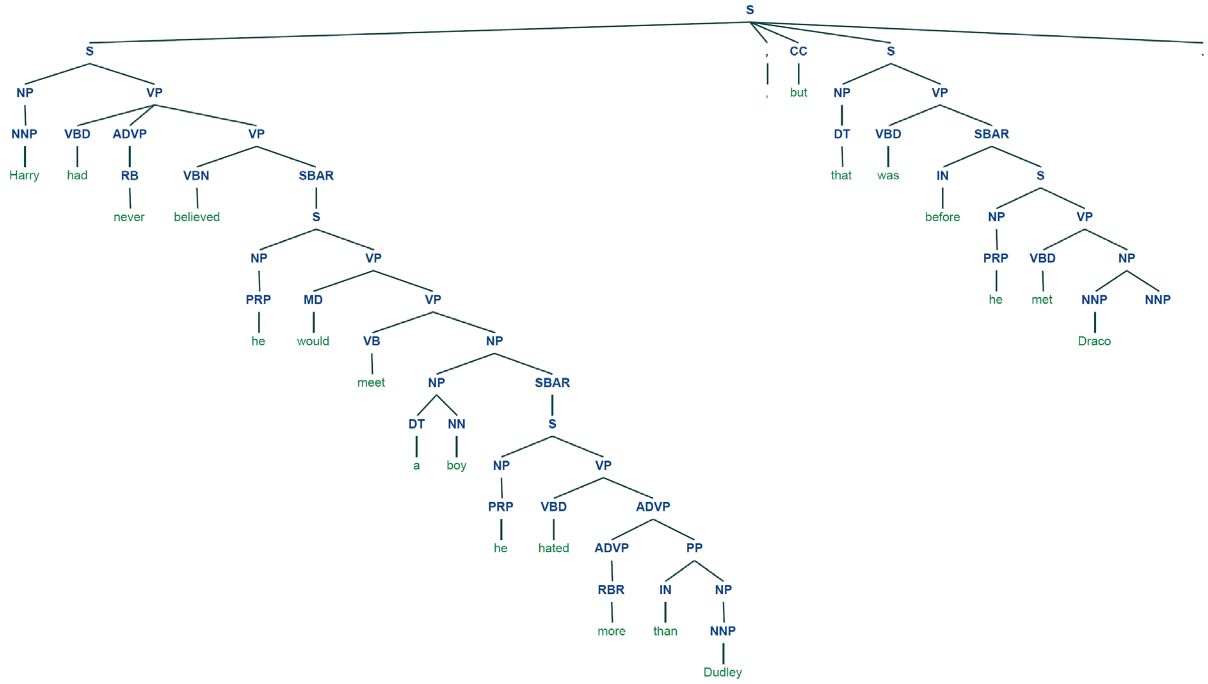

7. Malfoy (words seen till then = Harry had never believed he would meet a boy he hated more than Dudley, but that was before he met Draco Malfoy)

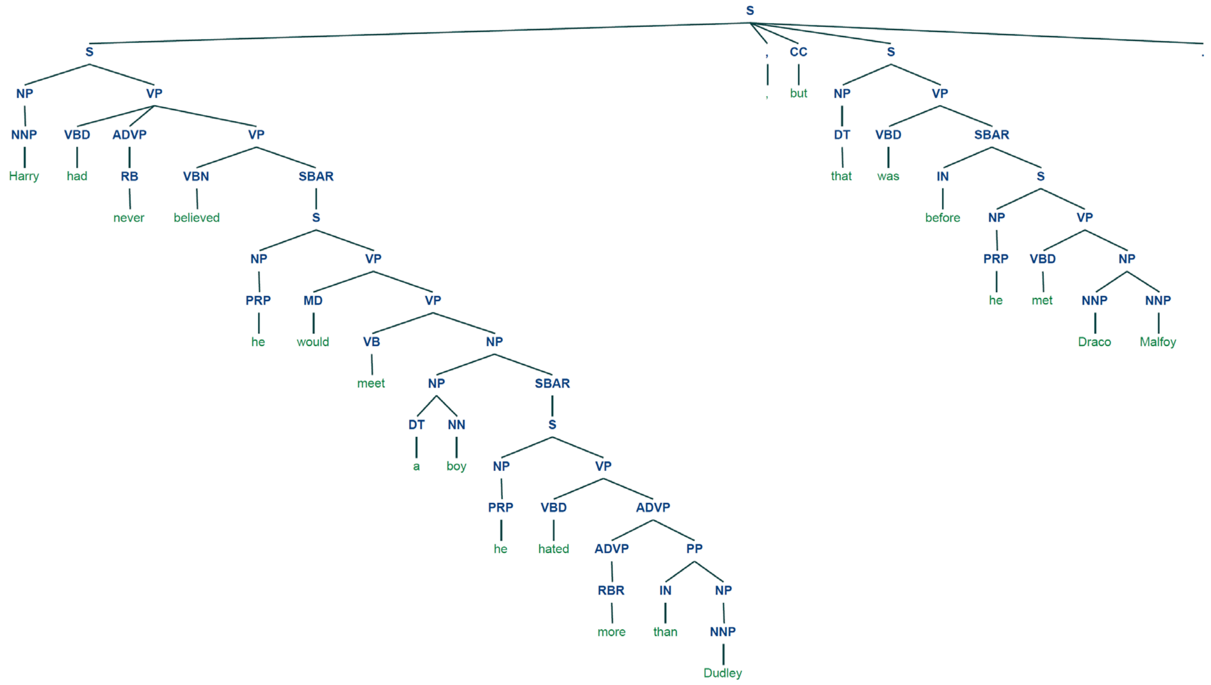


Again, each of these trees is encoded using the procedure described in the main paper to obtain one embedding per word.
